# Replicating Light-Off Startle Responses in *Drosophila melanogaster*

**DOI:** 10.1101/2021.02.24.432669

**Authors:** Thomas A. Allen, William J. Budenberg

## Abstract

We present a highly reproducible method for investigating the startle flight responses of wild type *Drosophila melanogaster* to light-off stimuli, using the automated Zantiks MWP unit. The built-in, live video-tracking of the Zantiks unit measured distance travelled between frames for 24 flies after light-off stimuli, whilst providing video-recordings of each startle. Using light-off stimuli which elicited peak startling, we found evidence for habituation of the startle response after only a few consecutive trials. Distance travelled on startle trials was reduced when a prepulse stimulus of shorter duration was introduced before the light-off stimulus, providing behavioural evidence for prepulse inhibition (PPI). Deficits in habituation and PPI are linked to various psychiatric disorders and our method holds great potential for use alongside genetic and pharmacological manipulations. Here, we demonstrate the capability of this highly automated, high throughput technology to streamline behavioural research on Drosophila, using a replicable, controlled environment.

## Introduction

Startle responses are crucial in an animal’s behavioural repertoire, facilitating evasion of predators and other threats. Such responses are exogenously provoked by salient stimuli with sudden onset (Eaton, 1984). For instance, rats startle when a loud noise reaches 80-90dB within 12ms of stimulus onset (Ladd, Plotsky and Davis, 2000). The neural mechanisms underlying startle responses vary between species but share a commonality in that they are fast-acting as to counteract imminent threats (Eaton, 1984). Consequently, reflex circuits are one of the primary mechanisms for implementing startle behaviour.

Whilst startle responses hold great ecological value, in an experimental setting adult *Drosophila melanogaster* have displayed startle responses to odour (Asztalos, Arora and Tully, 2007; Cho *et al*., 2004), and light-off stimuli (Fenckova *et al*., 2019). Using these stimuli, researchers have quantified habituation of *Drosophila* startle responses. The parameters constituting habituation are clearly defined by Thompson and Spencer (1966), outlining habituation as a reversible decrease in responding to a repeated stimulus. Startle habituation demonstrates an interaction between fast reflexive behaviour and slower learning, which helps prevent excessive responding to repetitive stimulus presentation. Moreover, Fenckova *et al*. (2019) established that impaired habituation of the light-off startle is related to *Drosophila* models of intellectual disability and Autism Spectrum disorder, suggesting translational applications for this body of research.

Our investigation primarily aimed to replicate the habituation of light-off startles observed by Fenckova *et al*. (2019) in a simple and reproducible fashion. We utilised the integrated, live video-tracking of the Zantiks MWP unit (https://zantiks.com/products/zantiks-mwp) to measure distance travelled by flies between frames. The unit provided a temperature-controlled, light-tight environment and allowed manipulation of light-off timing to the millisecond. All procedures were fully automated using scripts (Zantiks Ltd., 2021), enabling precise replication.

We first characterised the properties of light-off stimuli which elicit peak startling in wild-type adult *Drosophila*, considering colour and light-off duration. We then presented flies with consecutive light-off stimuli to assess habituation, using parameters closely resembling those used in Fenckova *et al*.’s paradigm. Initial characterisation of light-off stimuli revealed that short light-off durations induce startling to a lesser extent. We therefore investigated the potential of a short light-off duration stimulus to function as prepulse, inhibiting startle responses to longer light-off pulses. Prepulse inhibition (PPI) is a mechanism widely impacted in a range of neurological disorders, including schizophrenia, Huntington’s disease, and obsessive compulsive disorder (Fetcho and McLean, 2009), so warranted further examination.

## Methods and Materials

### Subjects

Subjects were wild type, adult *Drosophila melanogaster*, Oregon strain, sourced from Genetics Department, University of Cambridge. Flies were 3-10 days old of mixed sex. We tested a single plate of 24 flies in our light-off duration and habituation experiments. Three 24-well plates were used in our PPI experiment.

For each experiment, we transferred flies into wells using a brush, after cooling them on a chilled surface of cling film over ice. Wells were 16mm in diameter with 20mm spacing between wells. Wells were 6mm deep, with white walls and a clear acrylic base and lid. Approximately 2mm of solidified agar (2% agar, with 5% sugar and 1% nipagin, in water) covered the bottom of each well.

### Materials

We inserted well plates into the light-tight, experimental chamber of the Zantiks MWP unit, using the Zantiks LED light stimulation plate (see appendices). Light was dispersed by the insert such that light appeared from below the well plates. Light intensity, measured directly above the plate holder, was approximately 7000 lux.

Multiple Zantiks MWP units were used during testing. Allen, Harrison and Budenberg (2020) conducted work which established that differences in startle behaviour attributable to individual MWP unit and lab location were non-significant. Hence, we were able to run our PPI experiment on multiple units and different samples concurrently.

In a pilot study using 15ms light-off duration, qualitative examination of startle video-recordings revealed that green or white LEDs, opposed to red or blue LEDs, elicited the greatest startling. We used 530nm green LEDs in all subsequent investigations.

### Measures

The Zantiks MWP unit recorded movement of subjects using automated, live video-tracking. The unit recorded videos of each startle and provided a measurement of distance travelled by each fly in 1 second intervals. Startle flight responses were displayed in the video as rapid movements in straight lines, and hence startles were distinguishable from walking movements.

Through observation of startle responses in pilot videos, we found flight corresponded to long distances travelled in the 1 second, post-stimulus interval in the empirical data. Flight was rare in the absence of a light-off stimulus, with flies remaining still or walking. In the presence of the light-off stimulus, around two thirds of flies demonstrated the flight startle response, with flight relating to an approximate minimum of 20 pixels travelled in one second by an individual subject. By contrast, walking movements corresponded to 20 or fewer pixels travelled.

## Results

### Light-off duration

To assess the minimum light-off duration required to induce startle responses, we exposed flies to light-off stimuli with durations of 4ms, 8ms, 16ms, 32ms and 64ms. We ran 20 consecutive blocks of testing, in each of which durations were tested sequentially, from shortest to longest, with one minute between startles (figure 1).

**Figure 1.**
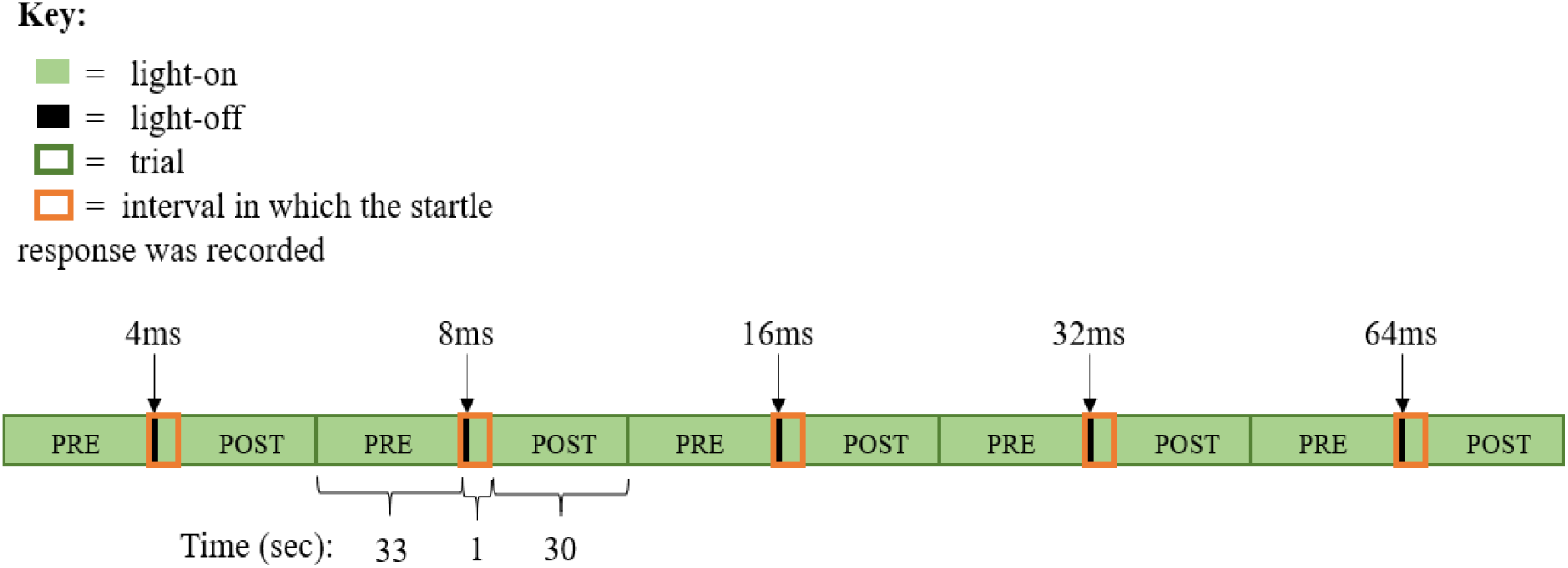
Procedural Outline of a Single Block Testing Light-Off Duration.

We found that 8ms light-off duration or longer was sufficient for inducing a startle response (figure 2). A one-way ANOVA on population distance travelled, with light-off duration as a within-subjects factor (4ms, 8ms, 16ms, 32ms, 64ms), revealed a significant effect of light-off duration on population distance travelled (F(4,95) = 37.9, p < .001). Post-hoc, multiple-comparison t-tests, with Bonferroni correction, revealed that each light-off duration condition between 8ms and 64ms inclusive differed significantly from the 4ms condition, but not from each other (figure 3). This is shown by significance markers in figure 2. Importantly, extending the light-off duration over 8ms did not significantly alter the distance travelled by the population.

**Figure 2.**
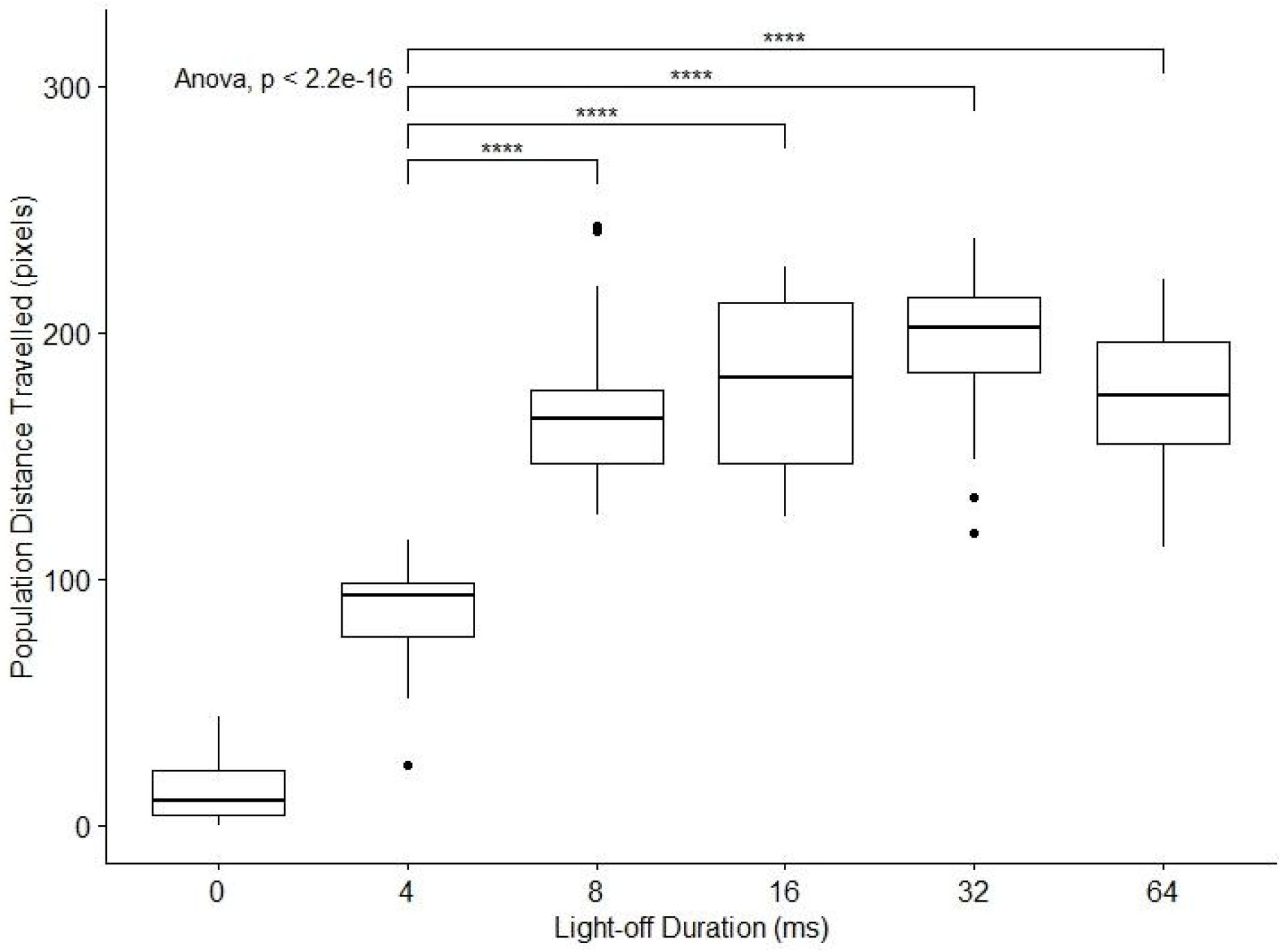
Population Distance Travelled in One Second Post-Stimulus for Each Light-off Duration. *Note.* Each box plot shows the quartiles and spread of population distances. Each datapoint is a trial, with 20 trials per light-off duration condition. 0ms light-off duration notates distance travelled in the 1 second, pre-stimulus interval across all 100 startle trials. Outliers, shown by dots, are 1.5 standard deviations below Q1 or above Q3.

**Figure 3.**
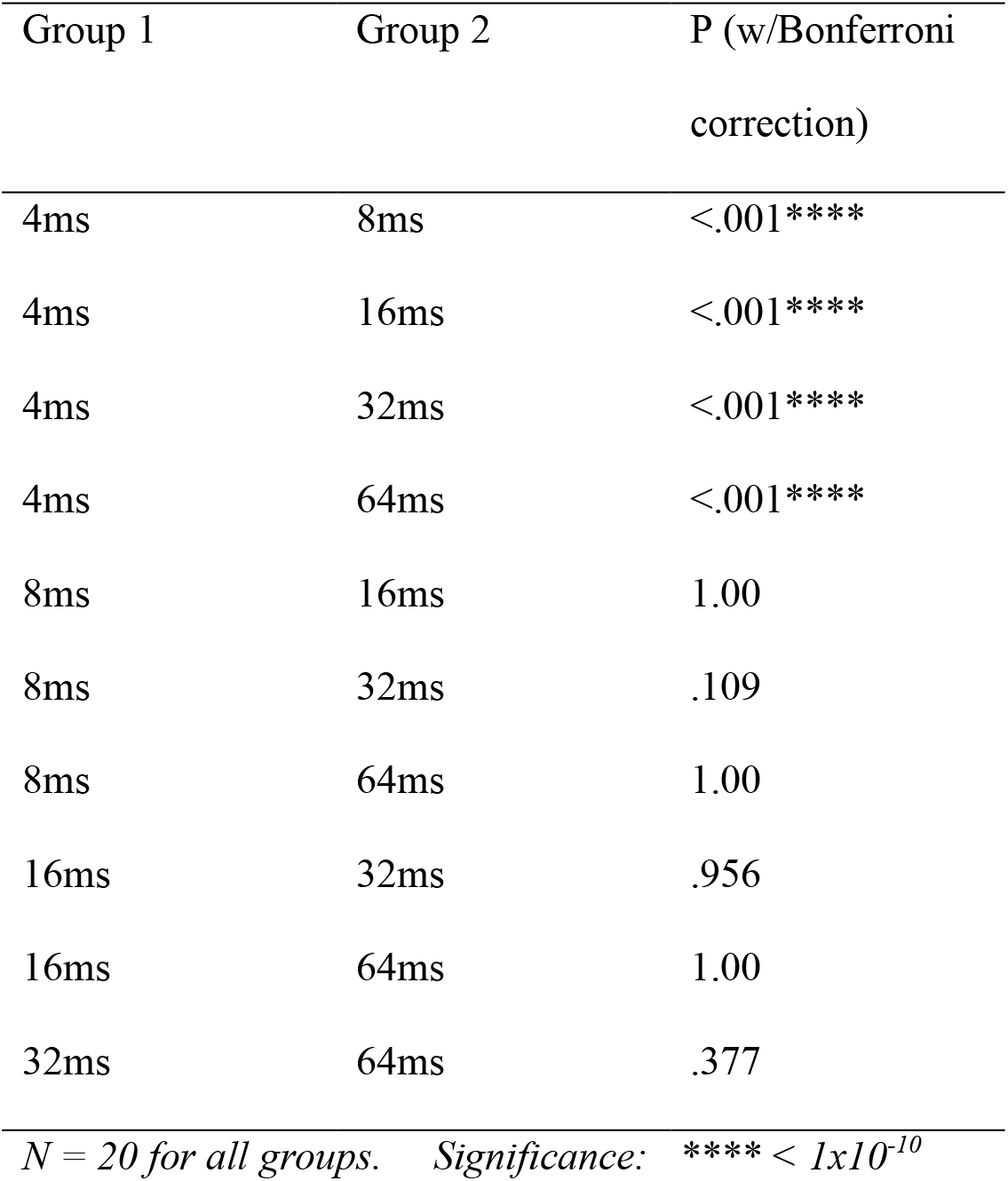
Multiple t-test Comparisons Results for Light-Off Duration Experiment.

### Habituation

In each block of our habituation experiment, we presented flies with 100 consecutive 15ms light-off stimuli, with a 1 second inter-trial interval. Whilst our light-off duration analysis confirmed that light-off stimuli over 8ms are appropriate, we selected these parameters to replicate the habituation effect observed by Fenckova *et al.* (2019). Three blocks of testing were conducted, with 60 minutes between blocks. Distance travelled was recorded during each one second trial.

Through graphical analysis, we found that startling diminished noticeably after only two consecutive trials, as depicted by figure 4. Our data is indicative of partial habituation to light-off stimuli, with an initial rapid decline in responding followed by noisy startling. Although small differences are visible between blocks, flies reliably startled most on the first trial in each block, demonstrating spontaneous recovery after 60 minutes. We did not conduct further statistical analysis on this dataset as we believe graphical analysis clearly demonstrates the patterns of responses.

**Figure 4.**
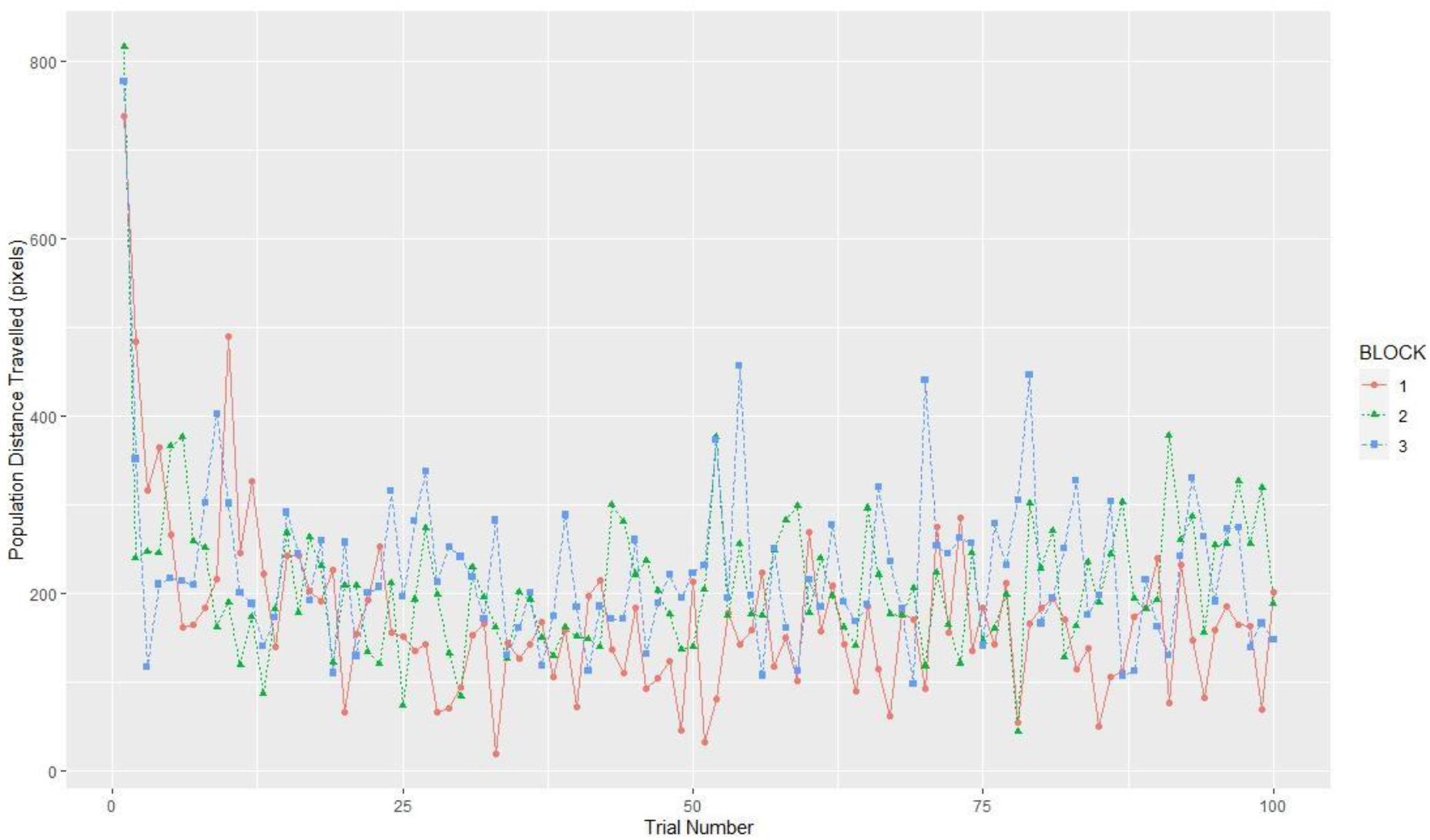
Habituation of Startles for the Population.

### Prepulse Inhibition

Investigation of the effect of light-off duration revealed that 4ms light-off stimuli elicit startles to a lesser extent than stimuli longer than 8ms. As such, short light-off stimuli could serve as prepulse stimuli which reduce responding to longer stimuli. Prepulse stimuli are typically less intense than their pulse counterparts but here we consider duration as equivalent to intensity, with longer duration comparable to a more intense stimulus.

Responses to a lone 15ms pulse were compared to a PPI condition in which the pulse was preceded by a 4ms prepulse (figure 5). The prepulse stimulus preceded the pulse by 30ms. A pilot study using variable prepulse-pulse latency (increasing by a factor of 2 from 15ms to a maximum of 960ms) suggested that this value was appropriate. We ran the experiment twice on three populations of 24 flies, totalling 6 sets of data. One of these 6 was excluded from analysis due to experimenter error in connecting the LED lights. In each run of the experiment, 16 PPI trials and 16 control trials were alternately tested, with an inter-trial interval of 20 seconds.

**Figure 5.**
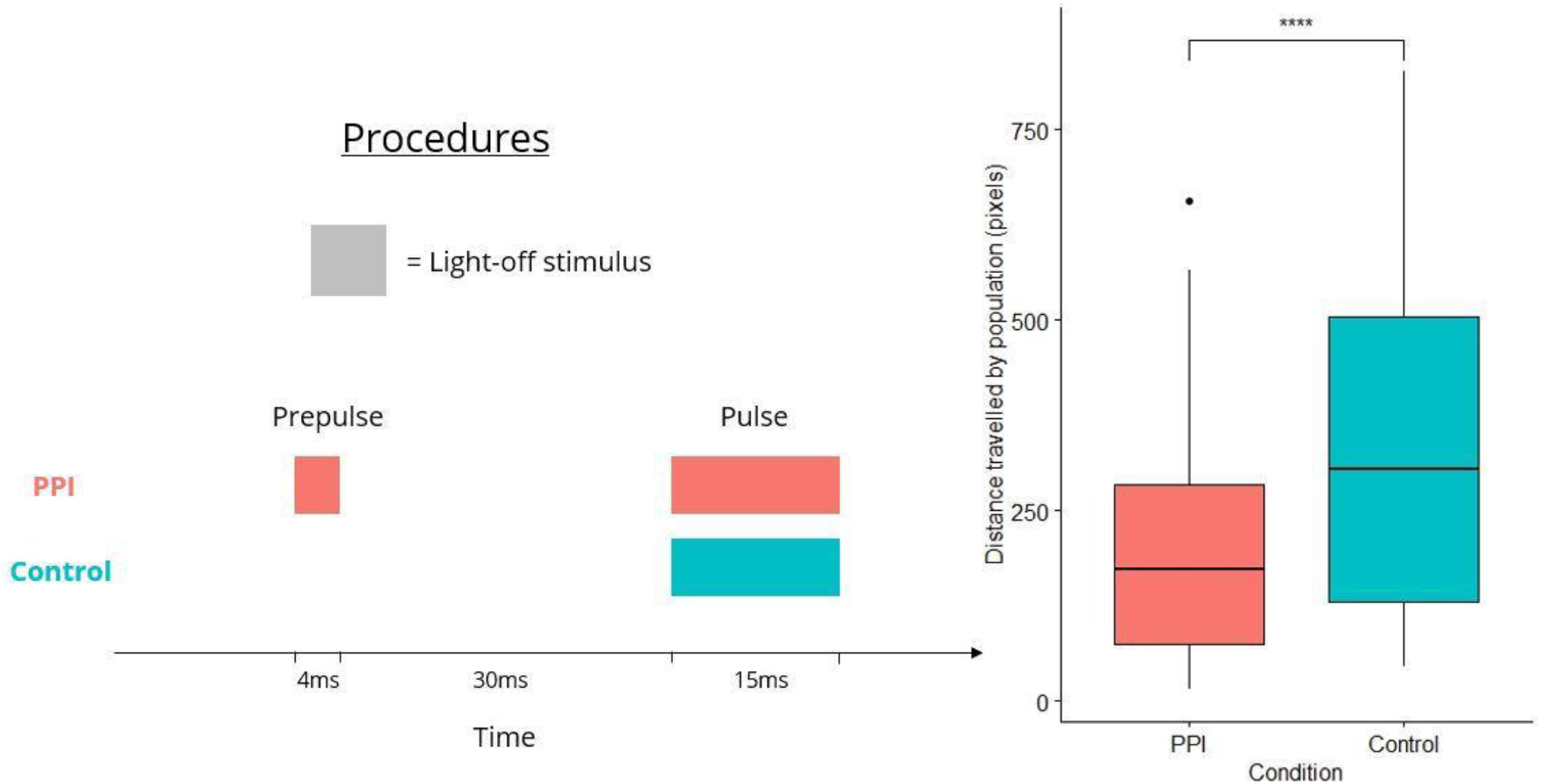
Procedure and Results of PPI Experiment. *Note.* The procedural outline (left) indicates the difference between PPI and control conditions. The graph (right) shows decreased startling during PPI trials as compared to control (pulse-only) trials.

Our results suggest that startling is decreased by the prepulse stimulus (figure 5). A paired samples t-test revealed a significant difference between population distance travelled in the PPI and control conditions (t(143) = 9.54, p < .001, Cohen’s d = .79). In both conditions, distance travelled was measured from the onset of the trial, rather than pulse stimulus onset.

## Discussion

We successfully reproduced behavioural evidence for habituation of the light-off startle response using our video-tracking method. Our results show a reversible decrease in responding with repeated stimulus presentation when using parameters matching those of Fenckova et al. (2019). Furthermore, we are the first to report PPI in adult *Drosophila*, finding that a short duration light-off stimulus can be used as a prepulse.

The success of our research should not only be measured by our findings, but also by the ease and accuracy with which our methods can be reproduced. Since we utilised a commercially-available behavioural unit, which carries out automated procedures, replication of our experiments is simple. In addition, the built-in environmental and stimulus control of the unit facilitates precise recreation of our paradigms. All procedural scripts and R data analysis scripts can be found on the Zantiks website (Zantiks Ltd., 2021).

With this in mind, we suggest that our method could be used in a deeper examination of light-off startle habituation. For instance, Rankin et al. (2009) define habituation as a response decrement ‘that does not involve sensory adaptation/sensory fatigue or motor fatigue’ but we did not rule out these alternative explanations for diminished responding. To distinguish between these accounts, a novel visual stimulus, to which adult *Drosophila* also startle, could be used to assess the stimulus-specificity of habituation within the visual modality. Failure to startle to the novel stimulus suggests either muscle fatigue, sensory fatigue, or sensory adaptation. In future, automated odour presentation may become a feature of the MWP unit which would enable the use of olfactory startle responses as a muscle fatigue test. Alternatively, demonstrating frequency-dependent spontaneous recovery, whereby shorter inter-trial intervals lead to faster spontaneous recovery, would also discredit the fatigue and adaptation accounts.

In spite of the clear patterns present in our behavioural data, there remains scope to fine-tune our measures. Although long distances correlate with taking flight during startles, distance travelled remains an indirect measurement, subject to noise from walking movements. To improve the signal-noise ratio in our data, we propose that a shorter window, of approximately 300ms (around 10 video frames at 30fps), be used in computing distance travelled. This would require no change in apparatus, only an alteration in our procedural script. Walking movements during this timeframe would be small in comparison to the large distance moved by startle flight. Then, distance travelled could be easily converted to a binary yes/no measure of startling by setting a minimum distance threshold, above which movement would be classified as a startle.

Nevertheless, our research has important implications in studying light-off startle responses of *Drosophila*. Whilst our investigations did not use genetic, toxicological, or neurological manipulations, we use basic principles for producing replicable research, applicable to a wide array of behavioural studies. We hope to have laid down groundwork for others to capitalise upon, with the potential to implement other methods in conjunction with the Zantiks MWP unit and examine more closely the underlying mechanisms of habituation and PPI in *Drosophila*. Adult *Drosophila melanogaster* may prove to be a useful model of intellectual disability and Autism spectrum disorder, as suggested by Fenckova et al. (2019), and of neurological disorders related to impaired PPI, such as schizophrenia.

*Drosophila* are by no means the only animal model of startle habituation and PPI. Zebrafish larvae, known to be a good genetic model of human disease (Howe et al., 2013), also startle to visual stimuli, showing habituation (Best et al., 2008) and PPI (Burgess and Granato, 2007). The Zantiks MWP unit’s video-tracking has been used to study startle behaviour in *Culex spp.* larvae (Harrison and Budenberg, 2021) and could be utilised in studying startle behaviour in a range of organisms. Moreover, larger Zantiks units, such as the LT, provide opportunity to run similar experimental procedures with model adult vertebrates, including mice, rats, fish, birds and xenopus.

Behavioural science has seen a shift in the past two decades towards a stronger focus on replicability. Difficulty in reproducing procedures and low powered designs both contribute to the ‘replicability crisis’ (Locey, 2020) and here we make a concerted effort to demonstrate the utility of the Zantiks MWP unit for easy replication of high powered, high throughput experiments. Other applications for the Zantiks units include being used to run automated batteries of behavioural phenotypic screening tests and to reliably reproduce tests in longitudinal studies across a population’s lifespan. As such, the technology we used here holds the potential to catalyse behavioural research, whilst upholding modern standards for easy and accurate replication.

## End Matter

## Author Contributions and Conflict of Interests

TAA and WJB designed and conducted the above experiments. TAA wrote the procedural scripts, data analysis scripts and the paper.

TAA carried out this work as part of a summer internship for Zantiks Ltd. WJB is the founder of Zantiks Ltd. and is an interested party in the use of Zantiks units.

## Acknowledgements

This study was entirely funded internally by Zantiks Ltd. as part of its research and development programme.

## Appendices

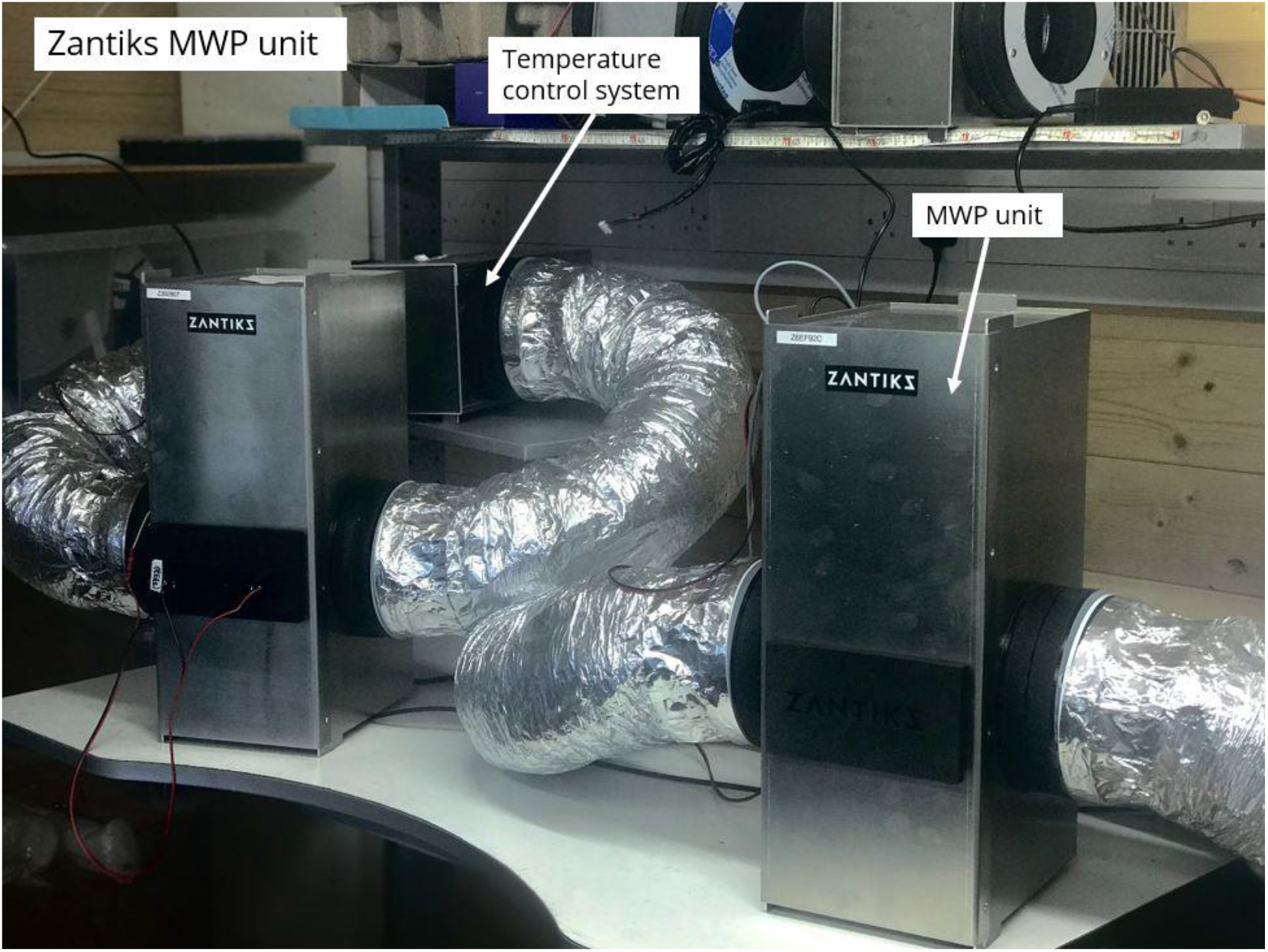

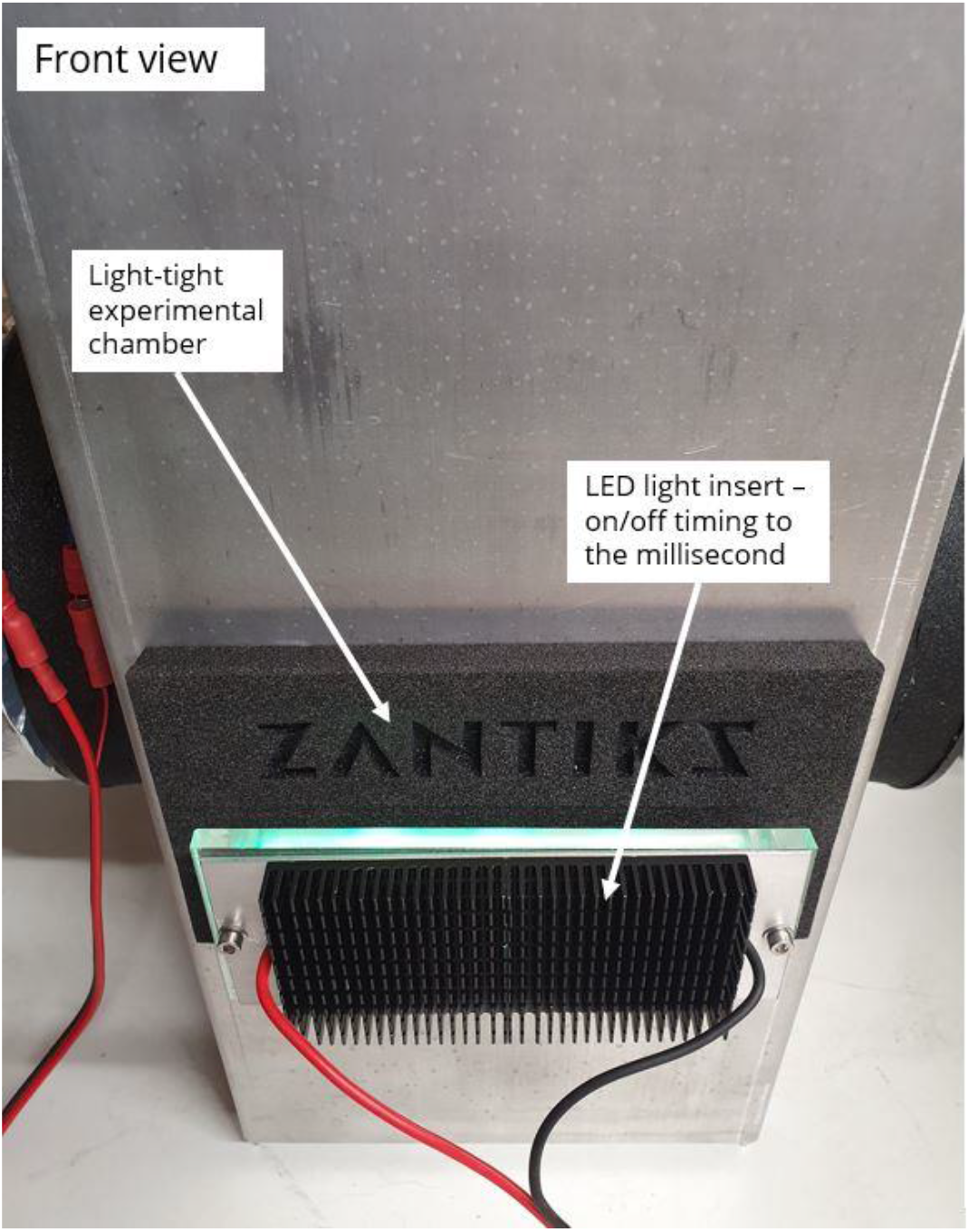

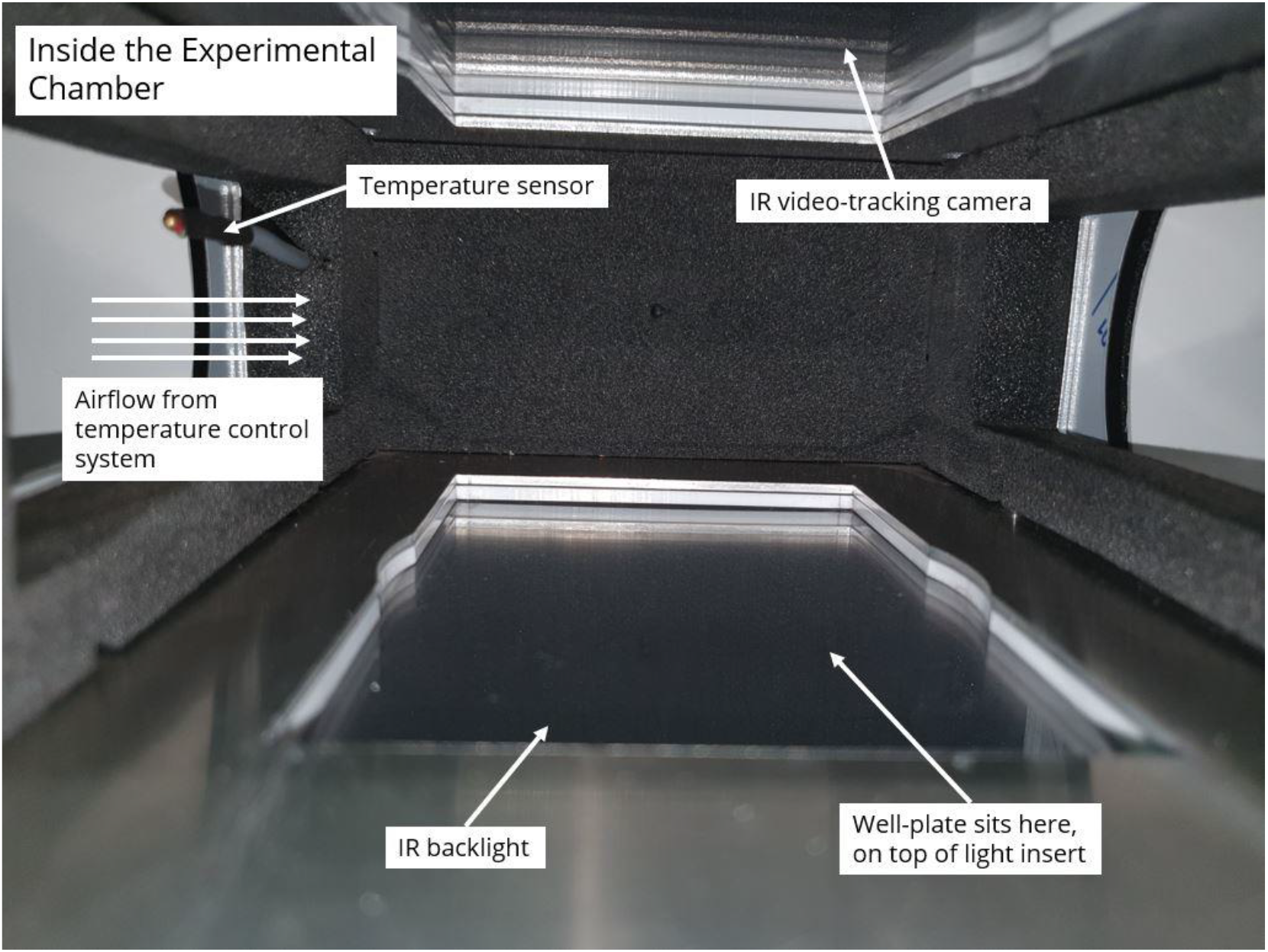

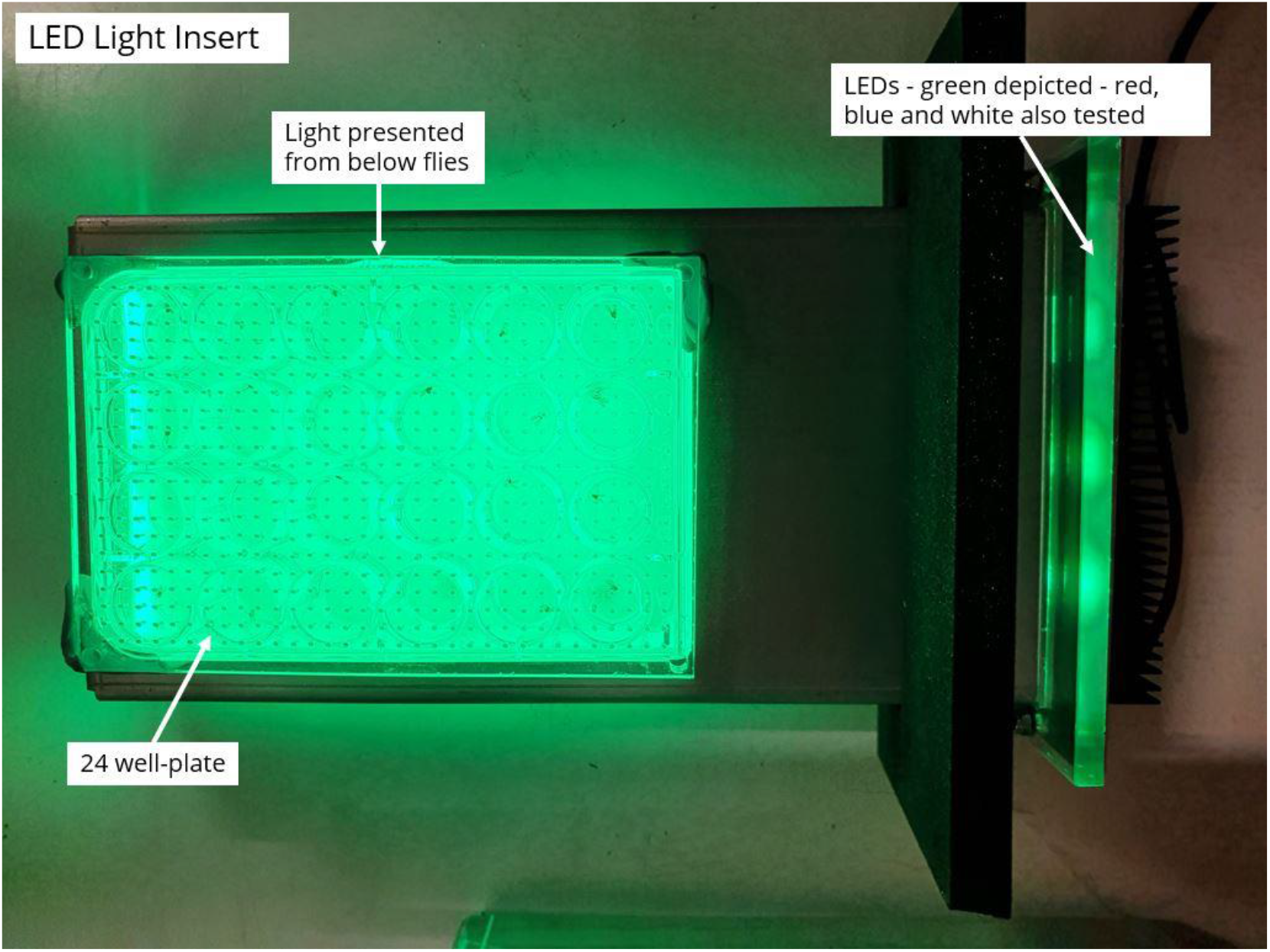

## Notes

https://zantiks.com/products/zantiks-mwp/

https://zantiks.com/support/script-library/script-library-template-1/

## References

Allen, T., Harrison E., & Budenberg W.J. (2020, November). Mosquito Larvae Reproducibility in Behavioural Science A Tale of Two Locations. [Conference poster]. Entomology 2020, Entomological Society of America virtual conference. DOI:10.13140/RG.2.2.10511.00161

Asztalos, Z., Arora, N., & Tully, T. (2007). Olfactory Jump Reflex Habituation in Drosophila and Effects of Classical Conditioning Mutations. Journal of Neurogenetics, 21(1–2), 1–18.

Best, J. D., Berghmans, S., Hunt, J. J. F. G., Clarke, S. C., Fleming, A., Goldsmith, P., & Roach, A. G. (2008). Non-Associative Learning in Larval Zebrafish. Neuropsychopharmacology, 33(5), 1206–1215.

Burgess, H. A., & Granato, M. (2007). Sensorimotor Gating in Larval Zebrafish. Journal of Neuroscience, 27(18), 4984–4994.

Cho, W., Heberlein, U., & Wolf, F. W. (2004). Habituation of an odorant-induced startle response in Drosophila. Genes, Brain, and Behavior, 3(3), 127–137.

Eaton, R. C. (1984). Neural mechanisms of startle behavior. Springer Science & Business Media.

Fenckova, M., Blok, L. E. R., Asztalos, L., Goodman, D. P., Cizek, P., Singgih, E. L., Glennon, J. C., IntHout, J., Zweier, C., Eichler, E. E., von Reyn, C. R., Bernier, R. A., Asztalos, Z., & Schenck, A. (2019). Habituation Learning Is a Widely Affected Mechanism in Drosophila Models of Intellectual Disability and Autism Spectrum Disorders. Biological Psychiatry, 86(4), 294–305.

Fetcho, J. R., & McLean, D. L. (2009). Startle response. In Encyclopedia of Neuroscience (pp. 375–379). Elsevier Ltd.

Harrison, E. R., & Budenberg, W. J. (2021). Mosquito Larvae (Culex spp.) Startle Responses to Vibration Stimuli. BioRxiv, 2021.02.18.431787.

Howe, K., Clark, M. D., Torroja, C. F., Torrance, J., Berthelot, C., Muffato, M., Collins, J. E., Humphray, S., McLaren, K., & Matthews, L. (2013). The zebrafish reference genome sequence and its relationship to the human genome. Nature, 496(7446), 498–503.

Ladd, C. O., Plotsky, P. M., & Davis, M. (2000). Startle response. George Fink. Encyclopedia of Stress.(Ed), 3.

Locey, M. L. (2020). The Evolution of Behavior Analysis: Toward a Replication Crisis? Perspectives on Behavior Science, 43(4), 655–675.

Rankin, C. H., Abrams, T., Barry, R. J., Bhatnagar, S., Clayton, D. F., Colombo, J., Coppola, G., Geyer, M. A., Glanzman, D. L., & Marsland, S. (2009). Habituation revisited: An updated and revised description of the behavioral characteristics of habituation. Neurobiology of Learning and Memory, 92(2), 135–138.

Thompson, R. F., & Spencer, W. A. (1966). Habituation: A model phenomenon for the study of neuronal substrates of behavior. Psychological Review, 73(1), 16.

Zantiks Ltd. (2021). Light-off Startle Response, (Adult Drosophila). https://zantiks.com/support/script-library/script-library-template-1

